# FEM-Based Mechanics Modeling of Bio-Inspired Compliant Mechanisms for Medical Applications

**DOI:** 10.1101/2020.06.15.151670

**Authors:** Yilun Sun, Dingzhi Zhang, Yuqing Liu, Tim C. Lueth

## Abstract

Compliant mechanisms are widely used in the design of medical robotics and devices because of their monolithic structure and high flexibility. Many compliant mechanisms derive their design ideas from nature, since the structure of biological organisms sometimes offers a better solution than the conventional mechanisms. However, the bio-inspired structures usually have very complex geometries which cannot be easily modeled and analyzed using traditional methods. In this paper, we present a novel finite element method (FEM) based modeling framework in Matlab to analyze the mechanics of different bio-inspired compliant mechanisms. Since the basic linear FEM formulation can only be employed to model small displacements of compliant mechanisms, a non-linear FEM formulation that integrates the modeling of large displacements, tendon-driven mechanisms and contact problems was implemented in the proposed framework to overcome the limitations. Simulations and experiments were also conducted to evaluate the performance of the modeling framework. Results have demonstrated the accuracy and plausibility of the proposed non-linear FEM formulation. Furthermore, the proposed framework can also be used to achieve structural optimization of bio-inspired compliant mechanisms.

## I. Introduction

IN the conventional medical devices or medical robotic systems, rigid-joint mechanisms are frequently used to realize motion transmission. Although the rigid links are stable and robust, they still have many limitations, such as complex assembly process and small gaps between joints, which make the sterilization of medical devices difficult and expensive [1]. To cope with these problems, compliant mechanisms are incorporated into medical device systems. Unlike the conventional rigid-joint mechanisms, the compliant mechanisms gain at least some of their mobility from the deflection of flexible members rather than only from movable joints [2]. With this interesting feature, it is possible to use a monolithic structure to realize medical devices that can perform complex tasks, which simplifies the assembly work and the sterilization process greatly. Since the flexible property of compliant mechanisms are quite close to the nature of biological organisms, many compliant mechanisms have a bio-inspired design [3]. For instance, the snake-like continuum robots get their biological inspiration from the flexible snake spines [4], [5]. Other examples include compliant medical forceps inspired by flytrap [6] and adaptive gripper inspired by fin-ray effect [7]. Fig. 1 shows the prototypes of the three mentioned bio-inspired compliant mechanisms, which are fabricated using different 3D-printers in our institute.

**Fig. 1.**
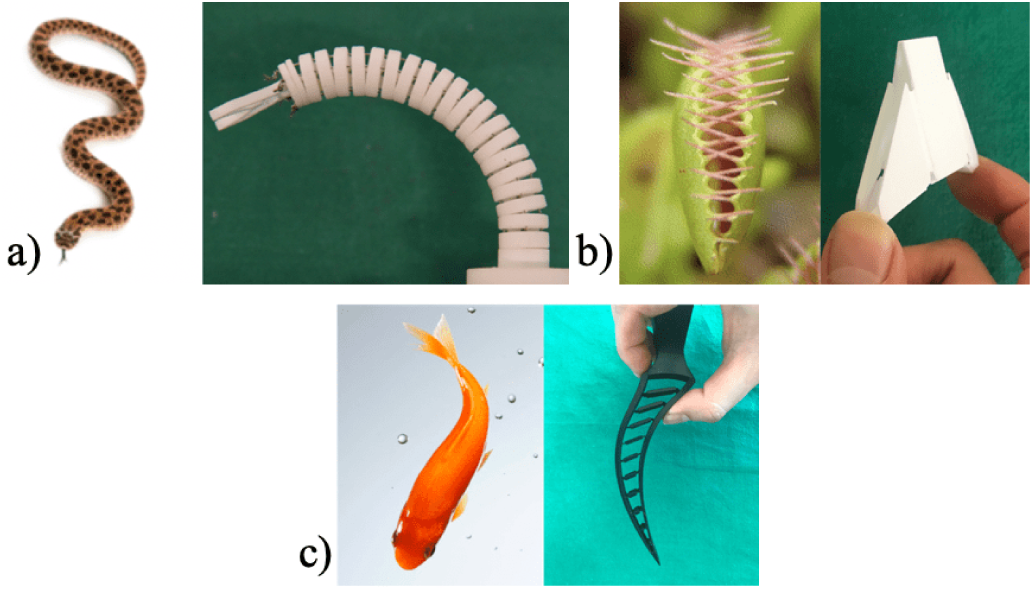
Examples of bio-inspired compliant mechanisms: a) A single-section snake-like soft robot, b) Flytrap-like compliant forceps, c) Finger of an adaptive gripper inspired by fin-ray effect.

However, due to their natural compliance, it is not easy to model and analyze the bio-inspired compliant mechanisms using the classic rigid-body mechanism theory, since the deformable continua cannot simply be treated as rigid bodies. To cope with this problem, a lot of methods have been developed to incorporate the elastic property into the kinematic analysis of bio-inspired compliant mechanisms. A popular method among them is the piecewise constant curvature model (PCCM), which is developed for analyzing snake-like tendon-driven continuum robots [8]. With the PCCM, each bent section of a snake-like soft robot, as is shown in Fig. 1a), is modeled as an arc with constant curvature. In [7], [9]–[11], various continuum robots inspired by snakes, elephant trunks and tendrils were successfully modeled and analyzed by the PCCM method. However, the PCCM method is mainly suitable for the cases of snake-like geometries, which cannot provide a general framework for the analysis of compliant mechanisms. Another important modeling method for compliant mechanisms is the pseudo-rigid-body-model (PRBM) method. With the PRBM method, the flexible parts with large deflections (the so-called flexure hinges) are simplified as a combination of torsional springs and rigid joints in a compliant mechanism [2]. In this way, the classic rigid-joint mechanism theory can also be used to analyze compliant mechanisms. In [12]–[14], the PRBM method was successfully employed to model continuum manipulators, compliant medical forceps and medical catheters. Although the PRBM method provides good results in calculating the final tip position of a compliant mechanism [12], the flexure hinges still have to be determined manually for model simplification, which makes the application of PRBM method for complex structures cumbersome. Another disadvantage of the PRBM method is that the detailed stress distribution in the compliant mechanism cannot be calculated due to model simplification. A more general modeling method for the compliant mechanism is the finite element method (FEM). With the FEM, the structure of a compliant mechanism is meshed into small elements. The detailed displacement and stress of each element can be then calculated based on the theory of continuum mechanics [15]. In this way, a compliant mechanism with any complex structure can be analyzed with a general framework as long as the structure can be discretized by a meshing algorithm, which is a great advantage over other modeling methods.

In the current state of the art, a lot of research work have incorporated FEM into the analysis of bio-inspired compliant mechanisms. In many of these work, the commercial FEM software were used to perform the analysis, as FEM has been an established method for several decades. In [16], Baek et al. have modeled the large deflections of concentric tube robots (CTR) using the FEM software ADINA. In this work, the structure of CTR was triangulated into shell elements, which is appropriate for the thin tubular structure but not applicable for general 3D structures. Different from [16], authors in [17] developed a modeling framework for soft material robots in Abaqus and Matlab which uses the general tetrahedral elements to mesh the structure. However, the proposed framework in [17] was mainly developed for analyzing snake-like soft robots, in which the FEM was employed only to extend the PCCM method. In [18], the software SolidWorks was used as a solid modeling and FEM tool to calculate the displacement and stress of a single section of a tendon-driven snake-like surgical robot. In this work, although the proposed FEM approach was more general, the load applied to the tendon-driven section was oversimplified as a force couple on the top of the section, which neglected the influence of the tensile force on the inner surface. To cope with these problems, authors in [19] have realized a general FEM framework for bio-inspired continuum manipulators, which has integrated the modeling of non-linear large displacements and tendon-driven mechanisms using self-implemented code. While producing more accurate finite element models than the previous work, the framework in [19] has still neglected some important features, such as the modeling of contact problems.

In this paper, we present a novel FEM-based modeling framework in Matlab to achieve the analysis of displacement and stress of various bio-inspired compliant mechanisms, such as snake-like continuum robots and bio-inspired compliant medical forceps. The entire framework was implemented in the Solid Geometry (SG) Library [20], a design platform that we developed in Matlab. The paper has the following contributions:

- A geometry modeling tool implemented in Matlab for creating 3D printable geometry models that can be meshed into tetrahedral elements.
- A non-linear FEM formulation of the mechanics of compliant mechanisms that integrates the modeling of large displacements, tendon-driven mechanisms and contact problems.
- The evaluation of the proposed modeling framework through its applications to the design and simulation of three different types of bio-inspired compliant mechanisms.

The remainder of the paper is organized as follows: Section II introduces the geometry modeling tool and explains the modeling approaches in the proposed non-linear FEM formulation. In Section III, applications of the modeling framework to three types of bio-inspired compliant mechanisms are presented. Section IV discusses the proposed modeling framework in terms of its performance and efficiency. Future work is also outlined. Conclusion is drawn in Section V.

## II. Modeling Methods

### A. Geometry Modeling Tool

There are two major representation schemata used in geometry modeling, which are Constructive Solid Geometry (CSG) and Boundary Representation (B-Rep) [21]. Although the CSG model is easy to implement, our geometry modeling tool employs the B-Rep principle since we focus on using 3D printing technology to fabricate bio-inspired compliant mechanisms and the standard data file for 3D printing is the B-Rep-based STL file. In the STL file, the geometry information is stored as a list of oriented triangles tessellating the surface of a 3D solid, which has a redundant storage of the vertex information and thus inefficient for data processing [22]. Therefore, we have employed the shared vertex data structure to implement the 3D geometry model, which is a struct in Matlab composed of an unrepeated Vertex List (VL) and a triangular Facet List (FL). On the other hand, since 3D solids are usually created by vertical and radial extrusion of 2D contours, we have also implemented the 2D contour model, which is a Matlab array containing the vertices along the contour boundary in a counterclockwise direction. For clearance, we named the 2D contour model and the 3D geometry model as Closed Polygon List (CPL) and Solid Geometry (SG), respectively. Fig. 2b) shows the SG of a flexure hinge, which is extruded vertically by the CPL in Fig. 2a).

**Fig. 2.**
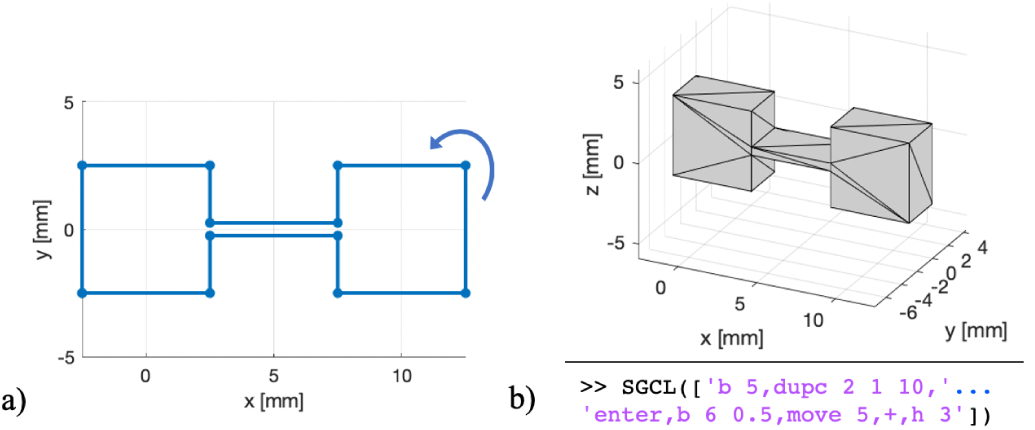
2D contour (CPL) and 3D geometry (SG) created by the geometry modeling tool: a) The CPL of a flexure hinge. The blue arrow indicates the listing directions of the vertices on the boundary, b) A SG extruded vertically by the CPL in Fig. 2a). The code in the figure shows the modeling process of the SG by using the SGCL modeling language.

Besides the data structure of the geometry model, we have also implemented many useful modeling functions in our tool, such as the creation of basic geometries and the manipulation of geometries through translation, rotation and Boolean operations. To simplify the modeling process of complex geometries, in which multiple modeling functions are involved, we have also developed a modeling language, the Solid Geometry Coding Language (SGCL). The SGCL language can encode the modeling steps of a geometry into a single string with an easy-to-learn syntax, and our modeling tool can interpret the coded string by calling the corresponding modeling functions. Table I provides an overview of some frequently used commands in the SGCL language. The code in Fig. 2b) has used the SGCL syntax to describe the modeling steps of the flexure hinge.

**TABLE I.** Commonly Used Commands of the SGCL Language

| Command | Illustration |
| --- | --- |
| b (x) (y) | Create a rectangular CPL with length $x$ and width $y$ |
| c (d) | Create a circular CPL with diameter $d$ |
| dupc ($n_x$) ($n_y$) ($l$) | Duplicate the current CPL for $n_x$ times in $x$ axis and $n_y$ times in $y$ axis, with a constant distance $l$ |
| h (z) | Create a SG by vertically extruding the current CPL with a certain height $z$ |
| move (x) (y) (z) | Translation of the current CPL or SG by a vector of (x,y,z) |
| enter | Put the current CPL or SG into stack and wait for another geometry for Boolean operation |
| +, -, & | Boolean operations (union, subtraction, intersection) of the current geometry and the geometry in the stack |
| rotx ($\theta$) | Rotation of the current SG along the $x$ axis by $\theta$ degree |

In this paper, we have employed Matlab’s Partial Differential Equation (PDE) Toolbox to mesh the created geometry model for the further FEM-based analysis. With the PDE Toolbox, a SG can be automatically and robustly meshed into a certain number of tetrahedral elements (see Fig. 3). The maximal and minimal element length can be adjusted by the user.

**Fig. 3.**
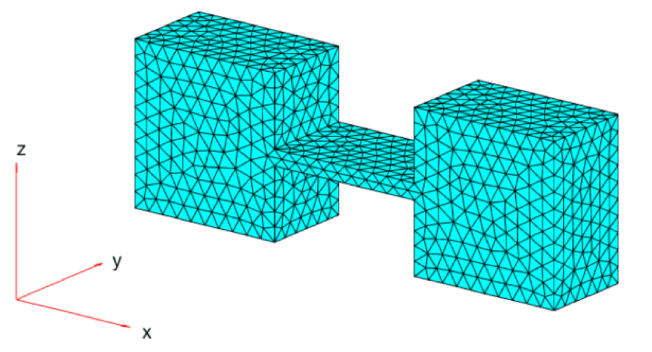
The meshed tetrahedral elements of the geometry in Fig. 2b) with the maximal element length of 0.5 mm.

### B. Modeling of Large Displacements

According to the linear FEM formulation for the static continuum mechanics [15], the displacement vector **U** of all nodes in an elastically deformed geometry can be calculated by solving the linear equation system in (1), where **K** and **F** are the global stiffness matrix and the external load vector, respectively. **KU** represents the internal force of the continuum and **K** is determined by assembling the stiffness matrix **K**_**e**_ of the *N* tetrahedral elements. The stress tensor ***σ*** and the strain tensor ***ε***(**U**) can be calculated by 2, where **E** and **B** are the elasticity tensor and the strain-displacement matrix, respectively.

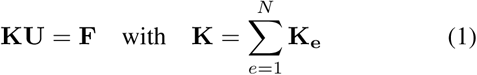

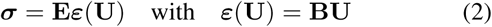

For the structure with small displacement **U**, the equation in (1) is reasonable and **K** can be treated as a constant matrix since the geometry changes little by the small displacement. However, for the large-displacement compliant mechanisms, **K** is inconstant and dependent on the displacement **U** during the large deflections. In this case, the equation system in (1) becomes a non-linear system which cannot be easily solved by the elimination method. To cope with this problem, we have developed an incremental-load method to integrate the large-displacement feature into our modeling framework. We firstly divide **F** equally into *N*_*inc*_ small increments **ΔF** and these incremental loads are applied to the continuum structure in *N*_*inc*_ steps. In this way, the linear FEM formulation in (1) can be easily modified as in (3) to approximate the incremental displacement Δ**U**_*i*_, since Δ**U**_*i*_ is so small that the linear theory is still valid. **K**_*i*_ is the global stiffness matrix of the deformed geometry in the *i*-th step. The total displacement **U**_*i*_ and stress tensor ***σ***_*i*_ can be obtained by accumulating Δ**U**_*i*_ and Δ***σ***_*i*_ as in (4) and (5), respectively.

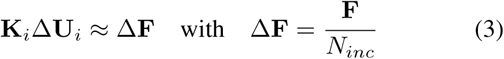

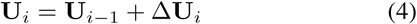

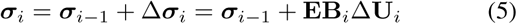

In this paper, we focus on modeling compliant mechanisms with incompressible materials. According to the law of mass balance, the volume of incompressible geometry should remain constant, even at large displacements. To achieve plausible simulation results, we have formulated the inequality in (6) to make the sum *ϵ*_*V*_ (*N*_*inc*_) of the volume errors smaller than a prescribed value *ϵ*_*V,tol*_.

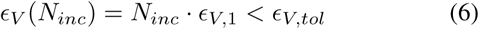

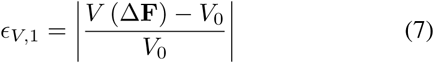

*ϵ*_*V*,1_ in (6) is illustrated by (7) that represents the volume error brought by the first incremental load, where *V*_0_ and *V* (Δ**F**) are the volume of the original and deformed geometry, respectively. *V* (Δ**F**) is calculated based on the updated coordinates of the geometry model with the linear displacement. Assuming that the volume error of each step is almost the same by applying the identical Δ**F**, *ϵ*_*V*_ (*N*_*inc*_) can be approximated as the product of *N*_*inc*_ and *ϵ*_*V*,1_. In this paper, we have employed the Matlab function `solve` to determine the minimum *N*_*inc*_ that satisfies (6). Algorithm 1 shows the implemented algorithm of the proposed incremental-load method.

#### Algorithm 1: Algorithm of the Incremental-Load Method for Modeling Large Displacements.

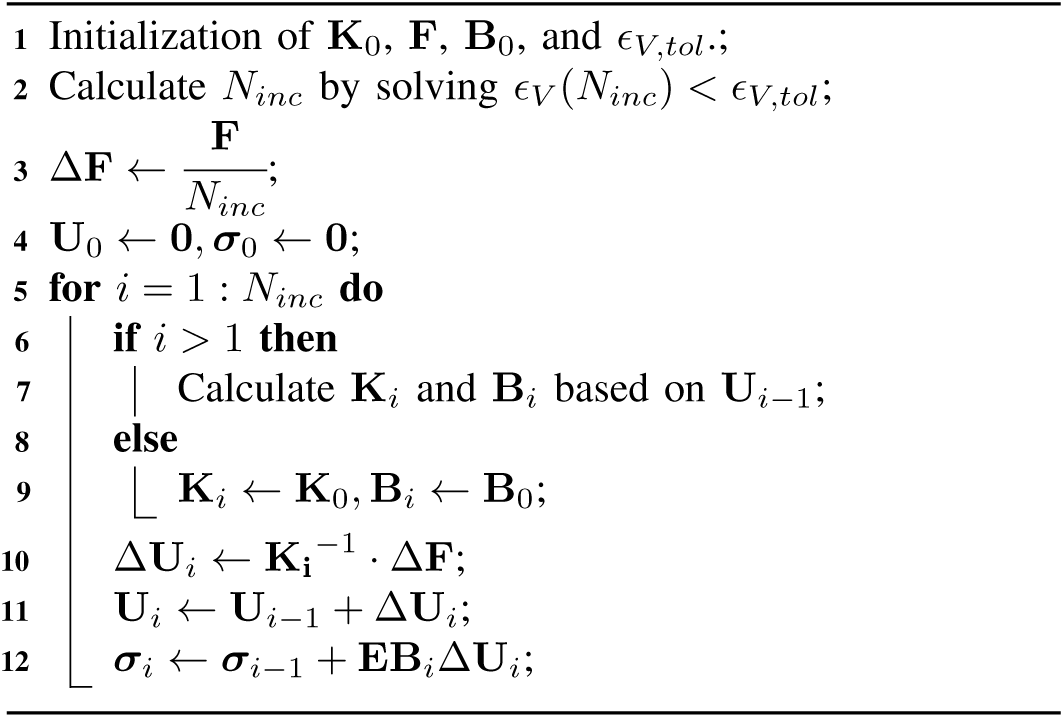

The large-displacement modeling of a compliant mechanism is graphically illustrated in Fig. 4, where the flexure hinge in Fig. 2b) is used as example and bent by a constant force. Firstly, user-defined geometries are created as fixed domains and loading domains for defining loading cases, as is shown in Fig. 4a). The nodes on the surface of the meshed geometry model, which are inside these user-defined domains, are detected for applying boundary conditions. Then, Algorithm 1 is used to calculate the large displacements. Fig. 4b) and c) show the modeling results by using the linear FEM (*N*_*inc*_ = 1) and the proposed incremental-load method ((*N*_*inc*_ *>* 1)), respectively. It can be noticed that, the linearly modeled geometry in Fig. 4b) swells greatly after one-step large displacement while the geometry volume of the non-linear model changes little during the incremental-load process. The advantage of the presented incremental-load method over the linear FEM is thus illustrated. On the other hand, the proposed method, although based on statics, can also be used to solve transient problems when the speed of movement is considered very slow. In our modeling framework, the incremental-load method is developed as the basis of the non-linear FEM formulation.

**Fig. 4.**
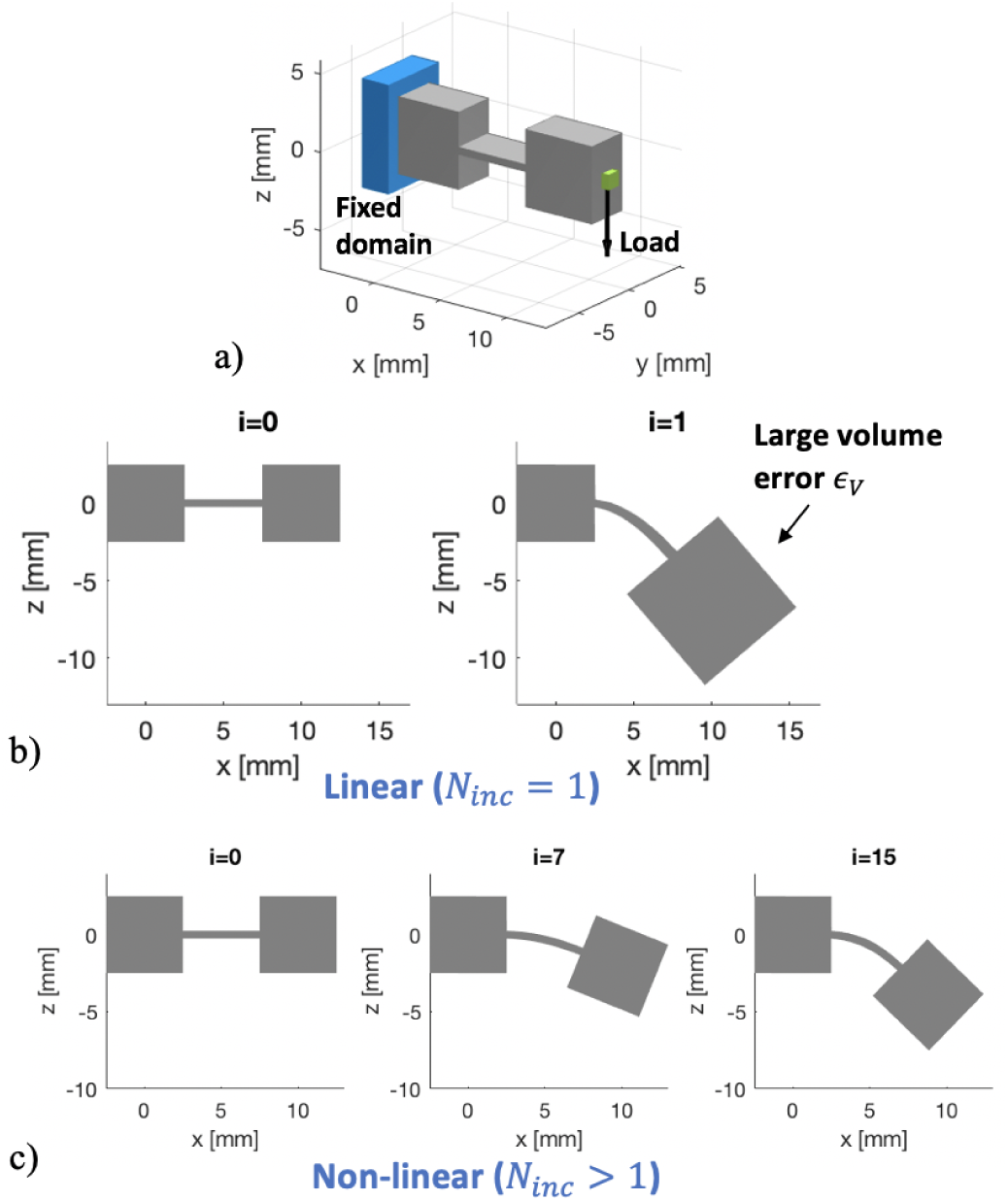
Large-displacement modeling of a compliant mechanism: a) Loading cases of a compliqnt mechanism (the flexure hinge in Fig. 2b) is used for illustration). The surface nodes in the green domain are loaded by a force in z-axis and the ones in the blue domain are fixed, b) The linear FE-analysis. The deformation is presented and a large volume error can be observed, c) The non-linear FE-analysis using the proposed incremental-load method. The intermediate shapes (*i* = 0, 7, 15) during the incremental-load process (*N*_*inc*_ = 15, *ϵ*_*V,tol*_ = 0.02) are presented, which show little volume error.

### C. Modeling of Tendon-Driven Mechanisms

Another important feature of our modeling framework is the modeling of tendon-driven mechanisms, as they are frequently used in the actuation of continuum robot systems, such as the snake-like soft robot in Fig. 1a). Fig. 5 shows the typical structure of a tendon-driven compliant mechanism, which is comprised of a bendable continuum body and a actuation tendon. The continuum body consists of a series of thin-wall flexure hinges and connecting disks, while the tendon is attached at the free end of the continuum body and goes through the disks to actuate the bendable structure. The tendon brings mainly two kinds of forces to the compliant mechanism, which are the tensile force *T* at the attaching point and the bending forces 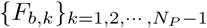 applied on the contact points of the disks, as is shown in the diagram of Fig. 5. The friction forces on the disks are not considered in this paper. In our notation, *b* stands for “bending” and *k* is the index of the *N*_*P*_ contact points ***P***_*k*_ between the tendon and the disks. As is graphically illustrated in Fig. 5, *F*_*b,k*_ can be treated as the addition of the tensile forces before and after the contact point ***P***_*k*_. As the tensile force in the entire tendon is identical, *F*_*b,k*_ can be calculated by using the following equations:

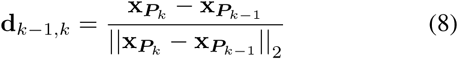

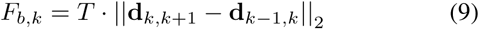

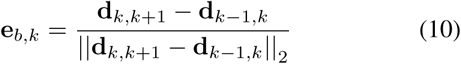

where 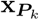 is the coordinate of ***P***_*k*_ and **d**_*k*−1,*k*_ indicates the normalized direction from ***P***_*k*−1_ to ***P***_*k*_. The force direction **e**_*b,k*_ of *F*_*b,k*_ can be determined by (10).

**Fig. 5.**
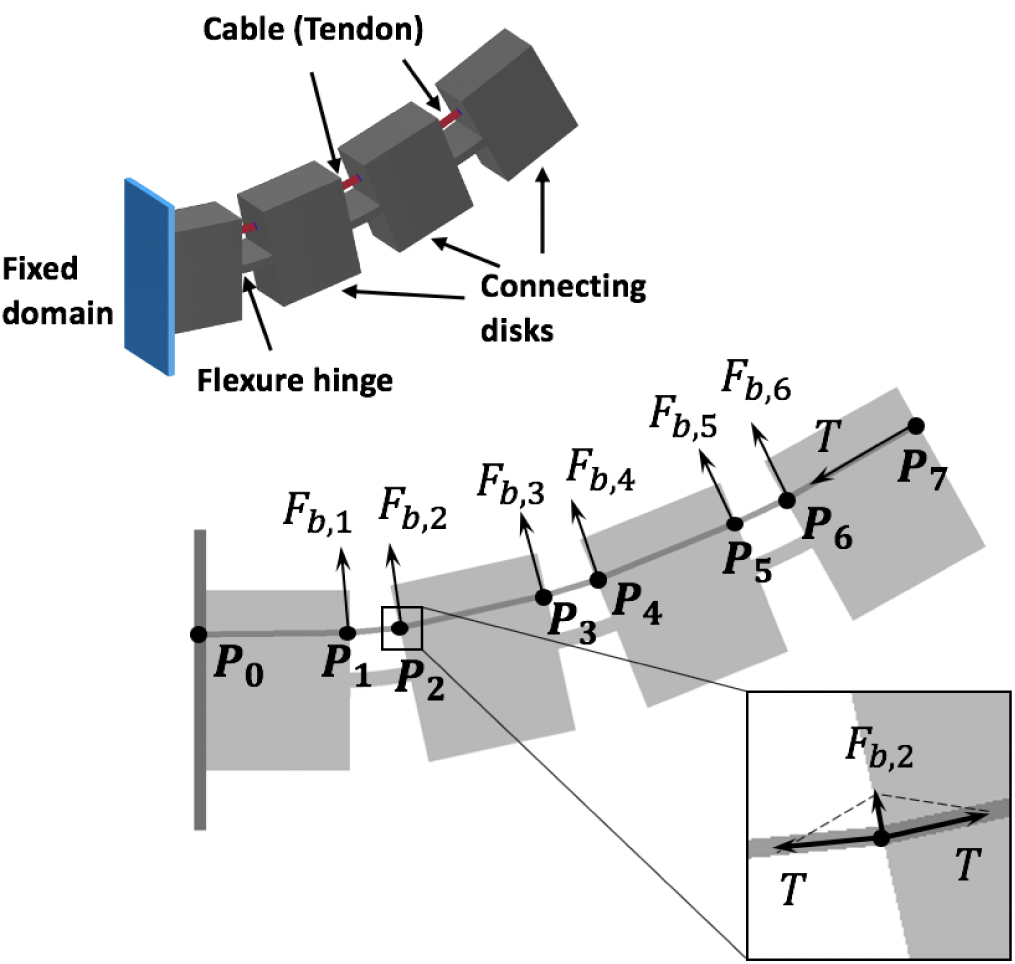
A compliant mechanism actuated by a single tendon. A schematic diagram is presented illustrating the principle of modeling the forces introduced by the tendon.

As can be seen in (9) and (10), the magnitude and direction of the forces introduced by the tendon are correlated with the displacement of the compliant mechanism, which introduces an additional non-linearity of **F** = **F**(**U**) to the FEM formulation. From this point of view, the incremental-load method in Section II-B cannot be directly used for modeling the tendon as the load vector **F** was assumed constant in Algorithm 1. To cope with this problem, we have extended the non-linear FEM formulation in Algorithm 1 to realize the modeling of tendon-driven mechanisms. Firstly, the tendon is defined as a series of loading domains as input for detecting the nodes of ***P***_*k*_. *T* is also predefined and incorporated into **F** for determining *N*_*inc*_ and the initial Δ**F** (similar to line 2 and 3 in Algorithm 1). Then, during the incremental-load process, the magnitude of the tendon force changes (Δ*T* and Δ*F*_*b,k*_) are calculated in each step by:

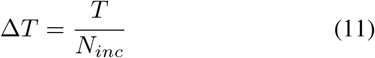

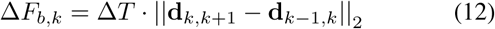

where the direction of Δ*T* and Δ*F*_*b,k*_ can be determined by (8) and (10) in each step, respectively. After that, we update the Δ**F** in line 10 of Algorithm 1 by accumulating the calculated Δ*T* and Δ*F*_*b,k*_ which are applied on the detected nodes of ***P***_*k*_. With these modifications in Algorithm 1, the non-linear feature of the tendon forces is integrated into our non-linear FEM formulation.

### D. Modeling of Contact Problems

As collisions between different parts often occur when a compliant mechanism undergoes large displacements, it is necessary to integrate the modeling of contact problems into our modeling framework for achieving plausible simulation results. The contact problems in continuum mechanics can be considered as a non-linearity of boundary conditions [23], which emerges when two or more surfaces penetrate each other. As can be seen in Fig. 6a), the contact-related non-linear boundary condition can be modeled as a pair of inverse forces applied on the contact boundaries and eliminating the penetration. To model the contact problems by using FEM, an important issue is to the detect the discrete nodes on the contact boundaries and then calculate the generated contact forces applied on the detected nodes. Herein, we have employed the slave-master concept and the node-to-facet contact search method from [23] for calculating contact forces. All contact problems in this paper are treated as frictionless.

**Fig. 6.**
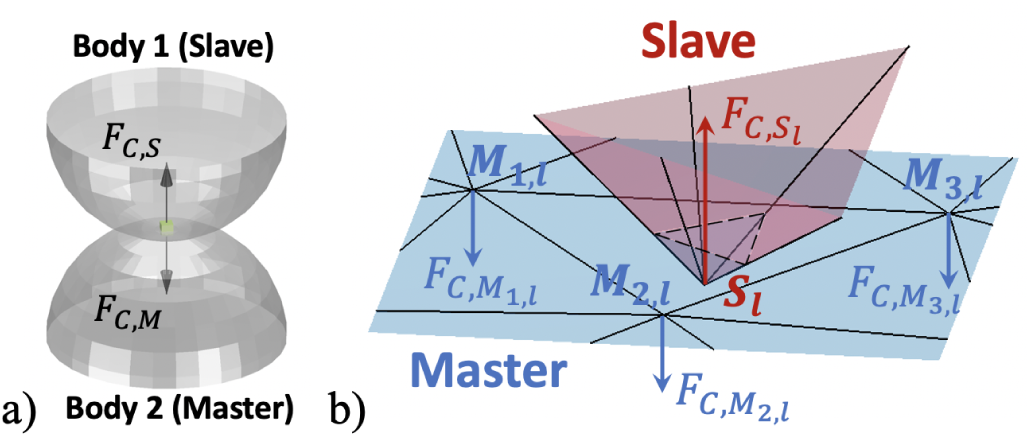
Graphical illustration of the modeling of contact problems: a) A typical contact problem of two continuum bodies. *F*_*C,S*_ and *F*_*C,M*_ are the generated contact force pair applied on the contact boundary (green domain), b) A diagram illustrating the modeling of contact forces using the slave-master concept and the node-to-facet contact search method.

The diagram in Fig. 6b) provides a graphical illustration of the node-to-facet contact search method. Firstly, a pair of geometries, called the slave body and the master body, are chosen as potential contact domains. Then, during the incremental-load process, all nodes on the slave surface are checked in each step to see if they penetrate any facet of the master surface. A contact problem is detected if any penetration between the slave nodes and the master facets occurs. Here, we suppose that a contact problem consists of *N*_*cp*_ contact pairs, where the *l*-th contact pair is comprised of a slave node ***S***_*l*_ and the corresponding master nodes of the penetrated facet {***M***_*j,l*_} _*j*=1,2,3_, as is shown in Fig. 6b). The contact forces generated by the *l*-th contact pair are calculated based on the principle that larger penetration needs larger contact force for elimination. Following this principle, the relationship between the slave contact force *F*_*C*,***S*** *l*_ and the penetration depth *g*_*l*_ is described by (15), where *K*_*C*_ is a scaling factor. *g*_*l*_ is defined in (14) as the normal distance between ***S***_*l*_ and the master facet 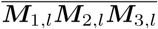, where **e**_*n,l*_ is the normal direction of 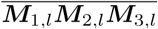. According to Newton’s third law of motion, the sum of the master contact forces is equal to *F*_*C*,***S*** *l*_ (see (16)).

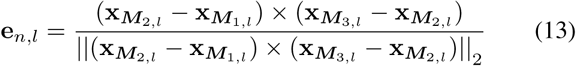

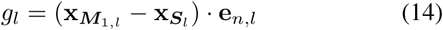

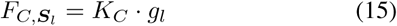

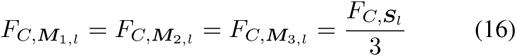

#### Algorithm 2: Algorithm of the Modeling of Contact Problems (Extending Line 10 in Algorithm 1).

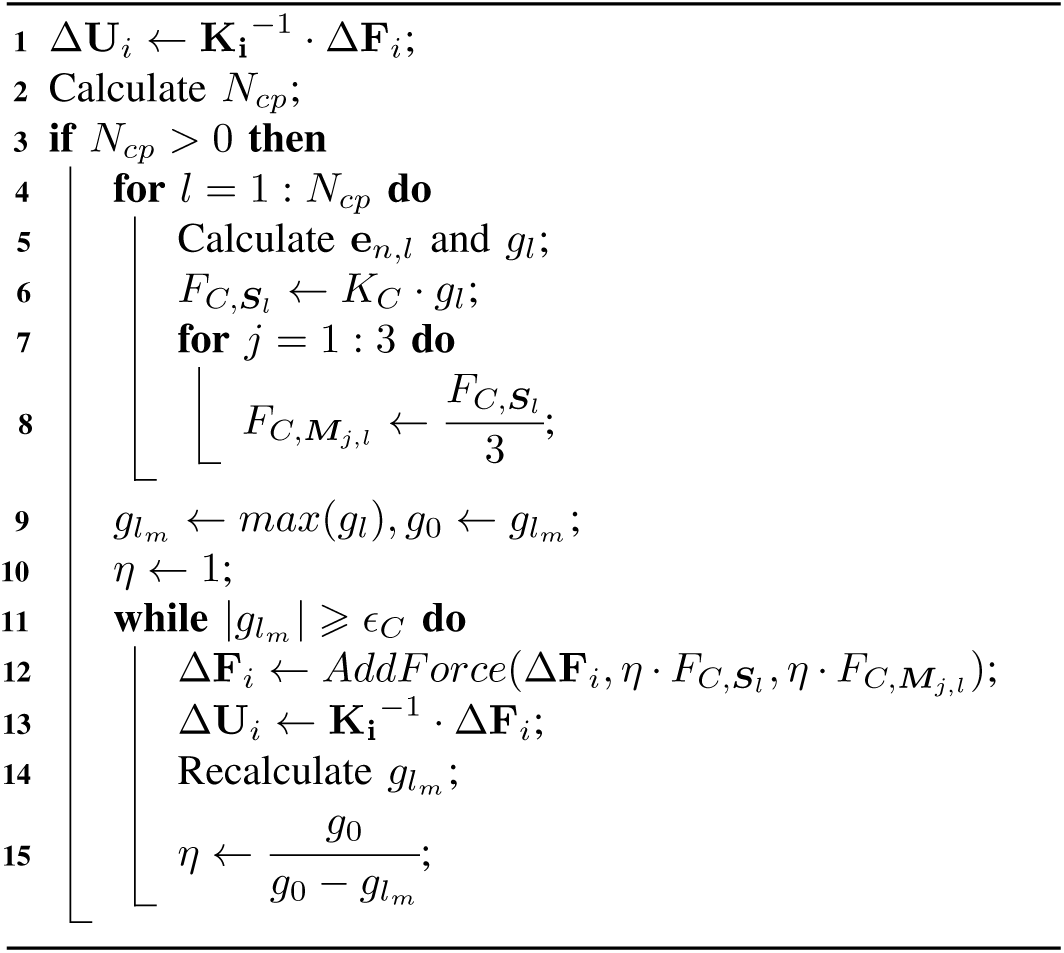

To ensure that the penetration can be successfully eliminated by the calculated contact forces, we have developed a numerical process to calculate *K*_*C*_, in which *K*_*C*_ is firstly assigned as the Young’s modulus *E* and then iteratively updated by multiplying the correction coefficient *η* in (17) until the absolute value of the maximum penetration depth 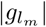 is smaller than a prescribed tolerance *ϵ*_*C*_ with the applied contact forces. *g*_0_ is the maximum penetration depth before applying the contact forces.

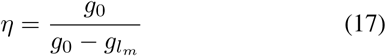

The presented modeling method of contact problems has been implemented in Algorithm 2, which can be integrated into our modeling framework by extending line 10 in Algorithm 1.

## III. Applications to Bio-Inspired Compliant Mechanisms

In this section, the proposed FEM-based modeling framework was applied in the design and simulation of three kinds of bio-inspired compliant mechanisms for medical applications. The simulation results were compared with the 3D-printed prototypes of the mechanisms to demonstrate the performance of the proposed modeling methods.

### A. Snake-Like Continuum Manipulator

In our institute, we are developing 3D printable robotic systems for minimally invasive surgery (MIS). The snake-like robots are used in our applications because of their high flexibility and tangible benefits to the patients. Fig. 7 shows the prototype of a selective laser sintered (SLS) continuum manipulator, which is equipped with a snake-like robotic structure for dexterous bending movements and an adaptive compliant forceps for safe manipulation of sensitive tissues [24]. Unlike the tendon-driven mechanism in Fig. 5, the orientations of the neighborin flexure hinges in Fig.7 are always vertical to each other and the continuum structure is actuated by two tendons. In this way, the continuum structure can be bent in different directions. In this section, we use the presented modeling framework to simulate the snake-like robotic structure actuated by a single tendon.

**Fig. 7.**
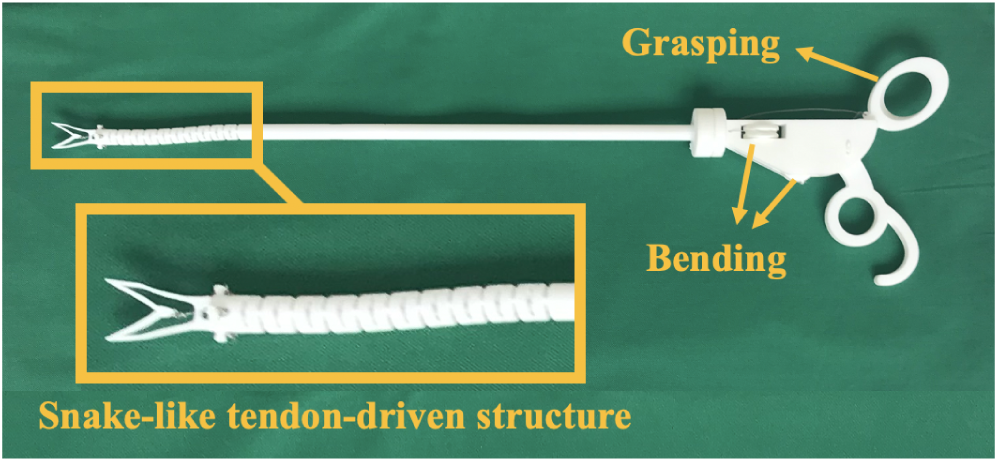
A SLS-printed continuum manipulator for robot-assisted MIS.

Fig. 8a) shows the geometry model of the snake-like structure, which is realized using the SGCL language. As the entire continuum manipulator is SLS-printed using polyamide (PA2200) [25] as material, the Young’s modulus and the Poisson’s ratio were set to 1700 MPa and 0.3 for calculating **K**. The maximum element size of the mesh was 0.5 mm. In the simulation, the bottom of the snake-like structure was fixed while a tendon was attached to the free end and went through the connecting disks. The intersecting domains between the tendon and the disks were detected for applying tendon forces. The contact-related master and slave surfaces were located using the cyan and green domains in Fig. 8 respectively to detect collisions between the disks. As the movement speed of the snake-like structure is considered slow in this paper, we have employed the proposed incremental-load method to analyze the transient bending process. In order to achieve a large bending angle, a large tensile force *T* of 5 N is applied in the simulation to actuate the snake-like structure. With a prescribed *ϵ*_*V,tol*_ of 0.02, the simulation was performed with 50 incremental steps and the results are presented in Fig. 8b). It can be noticed that, at the end of the simulation, the snake-like structure has achieved a large bending angle and the forceps of the manipulator has already passed the *xz*-plane of the coordinate system. From the simulated intermediate shapes we can see that, in the first part of the simulation (before the 30th step) the bent continuum structure had a constant curvature and was contact-free, while in the last simulation steps variable curvatures and collisions between the disks emerged, which shows the limitations of the PCCM method in the large bending angles. On the other hand, the stress distribution of the last simulation step, as is presented in Fig. 8b), indicates that the most stresses that emerged in the bent continuum body were concentrated in the flexure hinges near the fixed bottom, which also shows that the bending curvature was not identical in the entire structure.

**Fig. 8.**
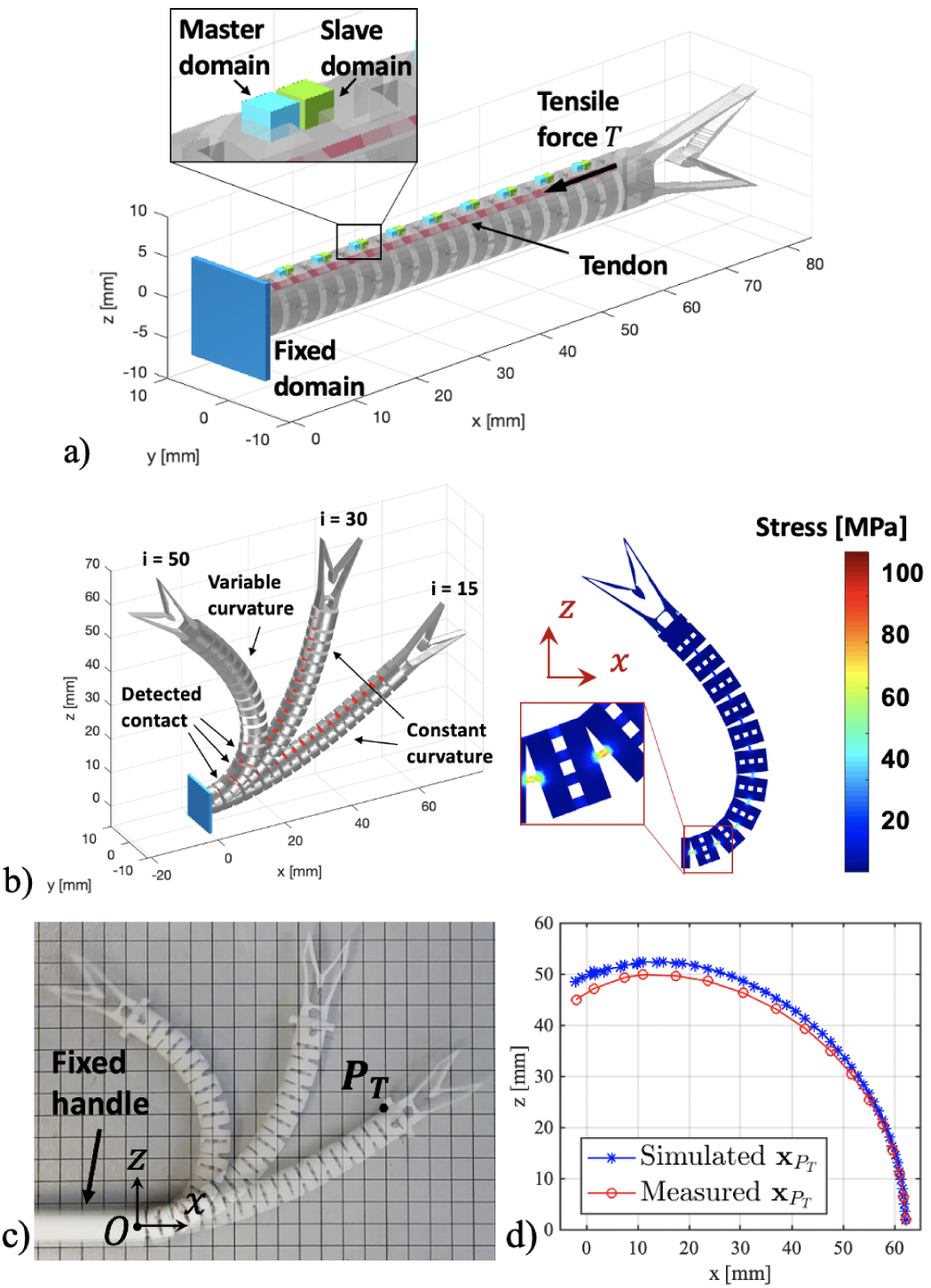
Modeling of the snake-like robotic structure in Fig. 7 and experimental evaluation: a) A schematic diagram showing the loading cases of the snake-like tendon-driven structure. The cyan and green domains are used to define the master and slave surfaces respectively, for detecting contact problems, b) Simulation of the snake-like structure. Intermediate shapes (*i* = 15, 30, 50) of the deformed tendon-driven mechanism and the final stress distribution are presented, c) Experiment for measuring the displacements of the snake-like structure, d) A comparison between the simulated (blue) and measured (red) coordinates of the attaching point ***P***_*T*_.

An experiment was also carried out to evaluate the plausibility of the simulated displacements. In the experiment, the handle of the continuum manipulator was fixed and the snake-like structure was constantly bent with the actuation of one tendon until the tendon attaching point ***P***_*T*_ passed the *z*-axis, as is shown in Fig. 8c). In this way, the trajectory in the experiment can be compared with that in the simulation. A digital microscope (Conrad DP-M17) [26] was used to measure the coordinates of ***P***_*T*_ during the bending movement. A comparison between the simulated and measured coordinates of ***P***_*T*_ is presented Fig. 8d). It can be noticed that, the simulated displacements were close to the reality, which shows the plausibility of our FEM-based modeling framework.

### B. Flytrap-Like Compliant Forceps

The second example of bio-inspired compliant mechanisms is a medical forceps inspired by Venus flytrap. The Venus flytrap is a small plant that catches insects with a trapping structure composed of two leaves (see Fig. 1b)). The trapping mechanism is realized by transforming the two leaves from a totally flat surface to a convex and a concave surface, which is highly flexible and efficient. Inspired by this interesting property, authors in [6] have developed a laser-cut compliant forceps with foldable origami structures, where the flexure hinges were fabricated using paper or metallic glass. Based on the work in [6], we have modeled the flytrap-like compliant forceps in this paper using our modeling framework and also fabricated it with the SLS printing technology. Since the polyamide has a high stiffness, the thickness of the folding area in the origami structure *t*_*fold*_ must be very thin so that the structure is bendable. To determine the optimal *t*_*fold*_, we have developed a structural optimization algorithm to iteratively modify *t*_*fold*_ so that the strain and stress inside the forceps are mainly concentrated in the folds. The algorithm can be illustrated by (18) and (19). The stress concentration factor *λ* of the folding area is defined as in (18), where 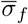 and 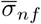 are the average stresses in the prescribed folding area and in the rest of the forceps, respectively. The updating scheme of *t*_*fold*_ is formulated as (19), where *t*_*fold,j*_ is the fold thickness in the *j*-th iteration while *λ*_0_ is the expected stress concentration factor. The optimization process stops when *λ* ⩽ *λ*_0_.

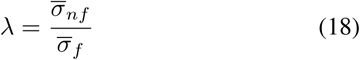

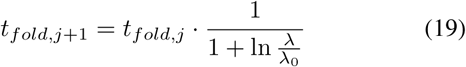

The modeling and optimization of the flytrap-like forceps is presented in Fig. 9. Fig. 9a) shows the realized basic geometry of the flytrap-like forceps. The maximum element size of the mesh was 1 mm in this case. In order to achieve the clamping movement, the center panel of the structure was fixed and forces were applied on the two side panels. The folds with the largest deformation, as they are marked in Fig. 9a), were selected for calculating 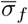. *t*_*fold*,0_ as set to 0.1 mm to start the optimization process. The optimization process converged at the 5th iteration and the result was presented in Fig. 9b). According to the calculated stress distribution in Fig. 9b), the highest stress was in the predefined folds, which is consistent with the optimization goal. Fig. 9c) shows the change of the stress distribution during the optimization process. With the optimized *t*_*fold*_, the SLS-printed flytrap-like forceps can successfully achieve the clamping movement, as is shown in Fig. 9d), which also shows the reliability our modeling framework.

**Fig. 9.**
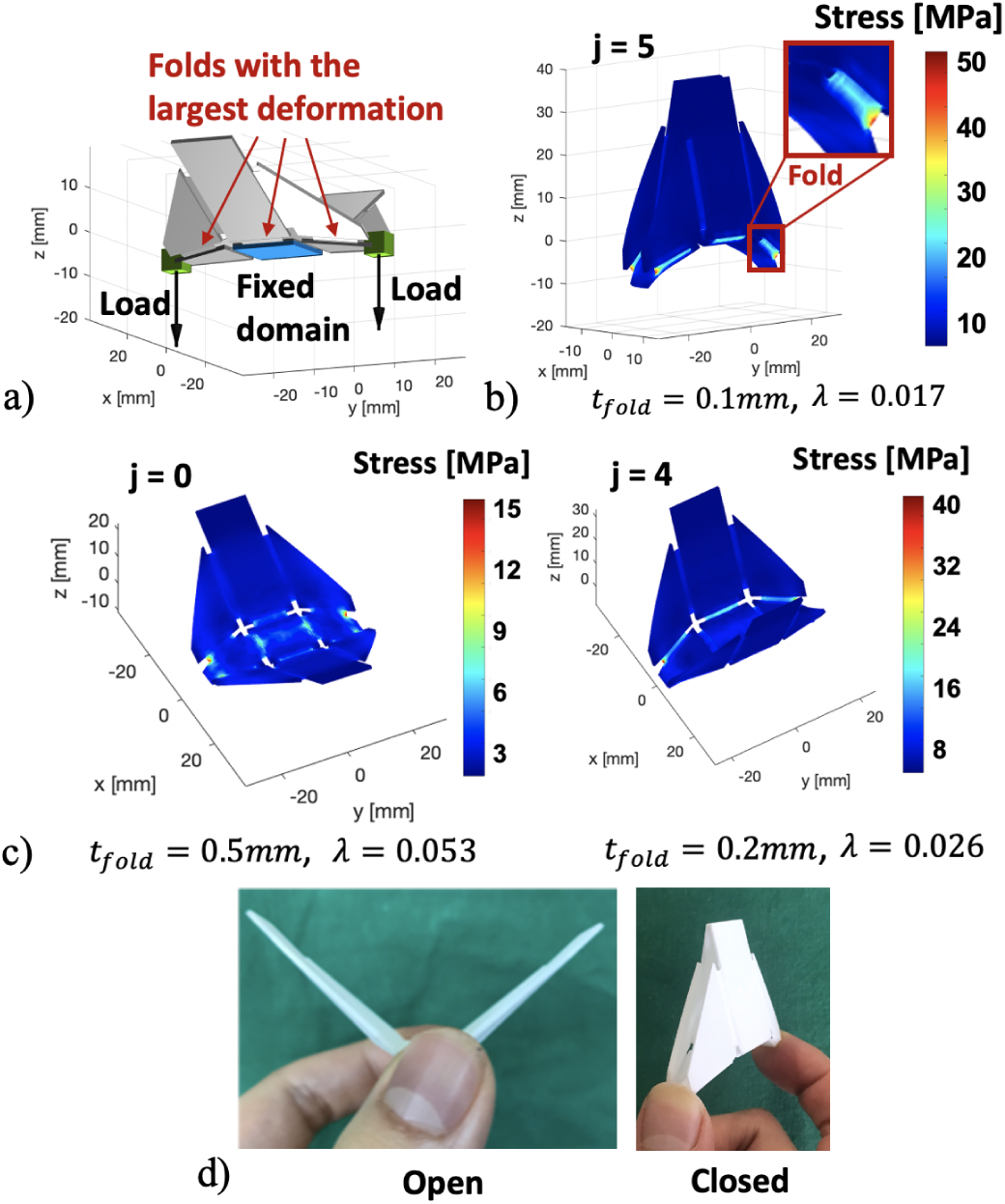
Modeling and structural optimization of the flytrap-like compliant forceps: a) A schematic diagram showing the loading cases of the flytrap-like compliant forceps, b) The calculated stress distribution of the optimized forceps, c) Stress distribution of the deformed forceps at some iterations of the optimization process, d) The SLS-printed forceps in the open and closed state.

### C. Fish-Fin-Inspired Adaptive Finger

The third example of bio-inspired compliant mechanisms is a fish-fin-like adaptive finger. As is shown in Fig. 1c), the biological fish fins are very flexible, allowing them to fit in almost any shape. Inspired by this interesting feature, authors in [7] have developed an adaptive compliant gripper, which has an isosceles triangular structure with multiple parallel stabilizers between its elastic sides. As the adaptive gripper could be used to safely grasp organs or tumors of irregular shapes (see Fig. 10a)), we have printed a finger of the adaptive gripper to explore its potential for medical applications. Since the polyamide is too stiff to realize such a flexible finger, we employed the stereolithography (SLA) printer (Formlabs Form2) [27] to fabricate the finger, using elastomer (Flexible Resin) [27] as material.

**Fig. 10.**
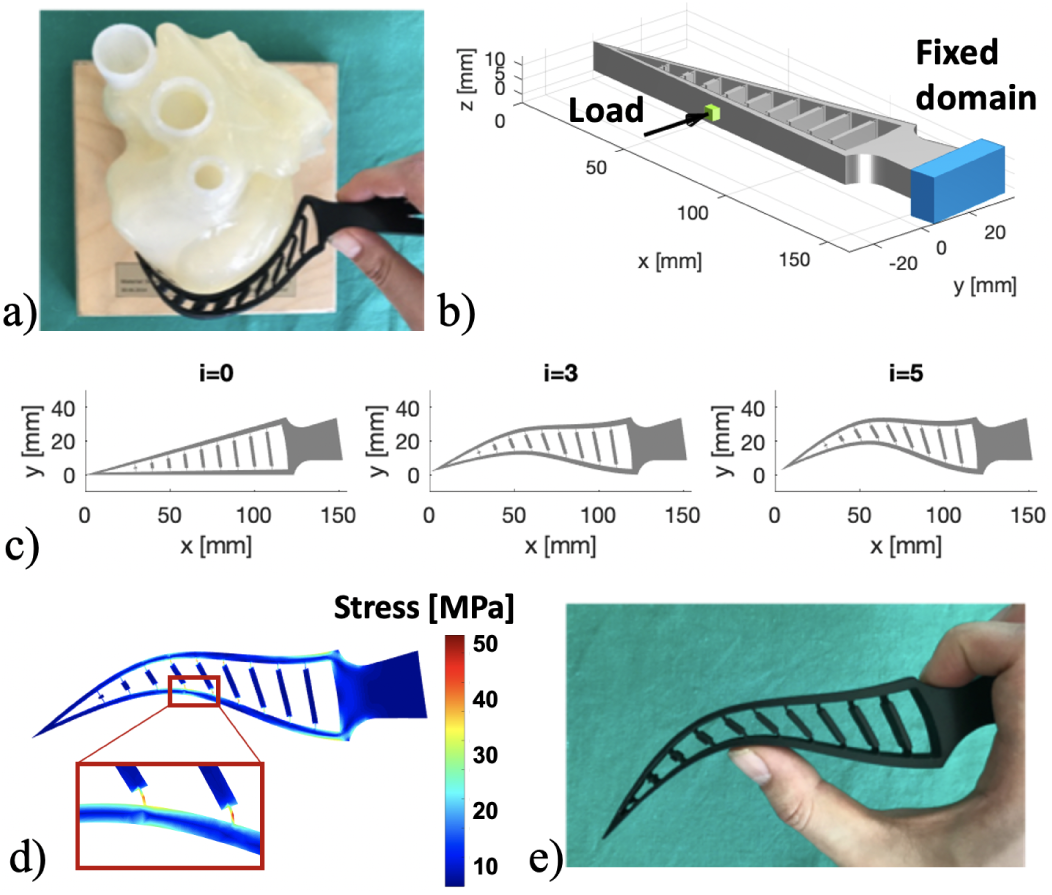
Modeling of the fish-fin-inspired adaptive finger: a) Using the adaptive finger to fit in the shape of a silicone heart model, b) A schematic diagram showing the loading cases of the adaptive finger, c) Simulation of the deformed adaptive finger. Intermediate deformed shapes (*i* = 0, 3, 5) are presented, d) The stress distribution of the deformed finger, e) The deformed shape of the SLA-printed finger.

Our FEM-based modeling framework was used to analyze the mechanical performance of the adaptive finger. Fig. 10b) shows the realized geometry of the finger. The Young’s modulus was set to 6 MPa according to [27]. In the simulation, a force of 0.5 N was applied on the middle part of the finger to mimic the effect of grasping an object, while the bottom of the finger was fixed. The simulation was performed with 5 incremental steps and the calculated intermediate shapes are presented in Fig. 10c). From the simulated stress distribution in Fig. 10d), it can be noticed that the highest stress was concentrated in the largely deformed flexure hinges between the parallel stabilizers and the elastic sides of the finger. From the biological point of view, this observation is useful for analyzing the actuation principle of the fin-ray effect. Besides, the calculated deformation of the last simulation step (*i* = 5) was close to the that of the deformed SLA-printed finger in Fig. 10e), which also shows that the proposed incremental-load method is plausible for modeling large-displacement compliant mechanisms.

## IV. Discussion

The work presented in this paper aims at analyzing the mechanics of bio-inspired compliant mechanisms using a general and accurate modeling framework. The proposed solution is based on FEM and can achieve general modeling of complex geometries. Our solution outperforms the previous work [9], [12], in which the PCCM and PRBM method are used for simplifying the geometries and the realized modeling frameworks are only applicable to the specific models. Besides, the non-linear FEM formulation in the proposed modeling framework has also integrated the modeling of contact problems which frequently occur in the bio-inspired compliant mechanisms, such as the snake-like tendon-driven structure, but are neglected in the previous work [19]. On the other hand, although many commercial FEM software can also perform robust FEM-based modeling of large-displacement compliant mechanisms, they cannot achieve the modeling of some specific mechanisms, such as the tendon-driven mechanism. In our framework, the tendon-driven mechanism has been integrated, as is presented in Section II-C. Another advantage of our modeling framework is that all modeling methods are implemented in Matlab and hence, additional interfaces between different platforms, as in the work [17], are no longer necessary, which greatly improves the efficiency of the framework. Furthermore, the example of the flytrap-like forceps in Section III-B has demonstrated the potential of our modeling framework for achieving structural optimization, which can be used to simplify the design process of the bio-inspired compliant mechanisms.

However, the framework can still be improved in several aspects. As is shown in Fig. 3, the PDE Toolbox always generates a mesh with a constant element size, which could make the simulation computationally expensive when the thin flexure hinges and the thick parts of the compliant structure have the same element size. This problem could be solved by introducing variable element size into the meshing algorithm of the PDE Toolbox, and will be further analyzed in the future. On the other hand, beside the displacement measurement, fatigue tests should also be carried out to evaluate the plausibility of the simulation results. In future work, we plan to integrate the modeling of other mechanisms, such as the pneumatic actuation, into the non-linear FEM formulation and deeply explore the potential of the proposed modeling framework for medical applications.

## V. Conclusion

In this paper, we proposed a novel FEM-based modeling framework in Matlab to analyze the mechanics of bio-inspired compliant mechanisms. The modeling framework is comprised of a geometry modeling tool and a non-linear FEM formulation that has integrated the modeling of large displacements, tendon-driven mechanisms and contact problems. Several modeling examples have been presented to show the performance of the proposed methods in modeling 3D-printed compliant medical robots and devices. Simulation and experimental results have demonstrated the accuracy and plausibility of the presented framework.

